# CCR4-NOT complex nuclease Caf1 is a novel shuttle factor involved in the degradation of ubiquitin-modified proteins by 26S proteasome

**DOI:** 10.1101/2020.05.13.093104

**Authors:** Ganapathi Kandasamy, Ashis Kumar Pradhan, R Palanimurugan

## Abstract

Protein degradation by ubiquitin proteasome system (UPS) is the major selective proteolytic pathway responsible for the degradation of short lived proteins ranging from regulatory proteins to abnormal proteins. Many diseases are associated with abnormal protein degradation; occasionally such dysregulated protein degradation is compensated by various transcriptional and translational control mechanisms in the cell. Among those pathways CCR4-NOT protein complex is responsible for transcriptional and transitional control of various gene expressions. Furthermore, CCR4-NOT complex also has a RING type ubiquitin ligase (E3) which is required for the degradation of several proteins. Here we report a novel function that the CCR4-NOT complex 3’-5’ exonuclease Caf1 is involved in ubiquitin-dependent degradation of short lived proteins by the 26S proteasome in yeast *Saccharomyces cerevisiae. caf1* deletion results in stabilization of R-Ura3 (N-end rule) and Ub-V^76^-Ura3 (Ubiquitin fusion degradation) substrates from proteasomal degradation. Additionally, *caf1* deletion accumulates ubiquitin-modified Ub-V^76^-Ura3 proteins and Caf1 binds to poly-ubiquitin conjugates and linear tetra ubiquitin chains. Surprisingly, Caf1 interacts with 19S regulatory particle complex of the 26S proteasome. Therefore, we conclude that Caf1 has an exciting novel function as an ubiquitin shuttle factor in which Caf1 targets ubiquitin-modified proteins to 26S proteasome for efficient degradation.

## Introduction

Timely degradation of both native and non-native proteins by Ubiquitin Proteasome System (UPS) and autophagy is an essential process for the cells to maintain proteostasis [1]. Proteolysis by 26S proteasome is a highly selective and tightly regulated process, which is accomplished by components of the ubiquitin-dependent and the ubiquitin-independent protein degradation pathways [1,2]. Ubiquitin-dependent proteolysis essentially requires the covalent attachment of poly-ubiquitin chains to substrate proteins and this process is carried out by protein ubiquitylation machineries involving cascade of enzymes E1 ubiquitin activating (Uba1), E2s ubiquitin conjugating (Ubcs) and E3s ubiquitin ligases [1]. The classical examples of such ubiquitin-dependent proteasomal substrates are UFD (Ubiquitin Fusion Degradation) and N-end rule [3–5]. These well studied substrate proteins harbour a degradation signal within them, for example, the UFD pathway substrate carries a non-cleavable ubiquitin attached at its N-terminal region, and the N-end rule substrates bears either type 1 (DERKH) or type 2 (LFWYI) amino acid residues as degradation signal (N-degrons) [6]. Cells are devoted with specific ubiquitylation machineries to attach polyubiquitin chains on substrate proteins, for example, the UFD pathway substrates are initially recognized and tagged with K29 linked ubiquitin chains by the E2 (Ubc4/5) and E3 (Ufd4), in the next step a heteromeric complex involving Cdc48^Ufd1/Npl4/Ufd2^ recognises ubiquitylated UFD substrates and further attaches K48-linked poly-ubiquitin chains that are responsible for proteasomal degradation [7,8]. Similarly, an ubiquitylation machinery complex involving Rad6 (E2) and Ubr1 (E3) recognizes the substrate proteins carrying N-degrons and attaches poly-ubiquitin chains [5,9]. Interestingly, a recent research work showed that ligases of the N-end rule and UFD proteolytic pathways Ubr1 and Ufd4 function as hetero-dimer complex to poly-ubiquitylate Mgt1 protein [9]. After substrate ubiquitylation step, a class of UBA-UBL domain containing proteins Rad23, Dsk2, Ddi1 binds to the polyubiquitylated substrate proteins via their UBA domain and delivers it to 26S proteasome by interacting with 19S regulatory particle complex via their UBL domain [10,11], there by these UBA-UBL proteins functions as ubiquitin shuttle factors during the proteolysis step. In parallel to the canonical ubiquitin shuttle factor proteins, in yeast the family of molecular chaperones Hsp70 and Hsp110 (Ssa1/2/3/4 and Sse1/2 in yeast) are reported in escorting polyubiquitylated proteins to 26S proteasome for proteolysis [12]. Interestingly, recent studies have shown that the ubiquitin ligases Ubr1 (involved in N-end rule pathway), Ufd4 (involved in UFD pathway) and Not4 (a stable CCR4-NOT complex subunit in yeast) are known to interact with 26S proteasome [13,14], however, the physiological significance of such interactions are currently unexplored.

CCR4-NOT is one of the largest multimeric protein complex that exist in all eukaryotes and functions in several cellular processes [15,16]. In yeast *Saccharomyces cerevisiae* the CCR4-NOT complex is composed of nine subunits [17]. Among those Not1 is the largest L-shaped protein that functions as a scaffold protein for assembly of remaining subunits namely Not2, Not3, Not4, Not5, Ccr4, Caf1, Caf40 and Caf130. Please refer the review [15] for extensive details about the composition on the CCR4-NOT complex in various organisms. Within the CCR4-NOT complex Ccr4 and Caf1 are the two 3’-5’ exonucleases and forms complex with Not1 which is responsible for de-adenylation of normal as well as defective mRNA [18]. The catalytic subunits of Ccr4 and Caf1 proteins are highly conserved among higher eukaryotes, but they differ in their deadenylase activity, for example, in yeast Ccr4 is the major mRNA deadenylation factor, but in mammalian cells Caf1 plays a major role [19]. Aside from being part of multimeric CCR4-NOT complex, the Ccr4 and Caf1 proteins forms independent subcomplexes separate from Not proteins [20,21]. In line with that, previous studies have reported that *caf1* deletion showed phenotypes similar to that of *ccr4* deletion in yeast [22]. Caf1 was recently shown to supress Htt103Q toxicity in yeast to ensure protein homeostasis [23,24] also pointing that Caf1 has widespread role in several cellular processes. Caf1 is known to be phosphorylated by Yak1p upon glucose limitation and the same study shows that glucose sensing is also mediated by Yak2 and Caf1(Pop2) pathway [25]. Intriguingly, it has been shown that mammalian CAF1 complements *pop2* deletion phenotype in yeast [26]. Another interesting observation shows that in *C. elegans* Caf1 is expressed both in embryos and in adults intriguingly the deletion of *C. elegans* caf1 (ccf1) results in embryonic and larval lethality [27] and in mammals Caf1 is essential for spermatogenesis [28]. Taken together Caf1 plays a vital role in higher eukaryotes. The remaining Caf40 and Caf130 proteins were identified as Ccr4 associated factors. The other subunits Not2, Not3, Not4 and Not5 docs the Not1 scaffold protein to form a stable CCR4-NOT complex [18]. Importantly, CCR4-NOT complex is implicated in several human diseases including diabetes, obesity and in liver associated diseases [16]. CCR4-NOT is a multifunctional protein complex having function in chromatin remodelling, transcription elongation, mRNA transport, mRNA decapping, mRNA deadenylation and mRNA quality control. CCR4-NOT complex proteins were found to be associated with chromatin remodelling proteins, RNA polymerase, ribosome, proteasome and molecular chaperones [15,29–32]. The interaction of CCR4-NOT complex subunits Ccr4, Not4 and Not1 with 26S proteasome and molecular chaperones supports the connection of CCR4-NOT in protein degradation pathways. In line with that the RING type ubiquitin ligase Not4 has been reported in the proteolysis of translationally arrested peptides and misfolded proteins [22,33,34], however, the Not4 catalytic RING domain was shown to be dispensable for such function. In addition, Not4 has so far been known to monoubiquitylate Rpt5 subunit of the 26S proteasome *in vitro* [35], furthermore, the substrate at which Not4 polyubiquitylates remains to be explored. Besides, Not4 the function of other CCR4-NOT complex proteins Caf130, Caf40, Caf1, and Not2 in the UPS pathway is totally unexplored. Therefore, in the current study we systematically tested the requirement of CCR4-NOT complex subunits in the protein degradation pathway. We discovered that the Ccr4 associated factor Caf1/ Pop2 has a novel function in the UPS pathway, Caf1 is specifically required for the ubiquitin-dependent degradation of N-end rule and UFD pathway substrates. *caf1* deletion results in accumulation of UFD substrate in the poly-ubiquitylated forms. Interestingly, we found that Caf1 interacts with cellular ubiquitin conjugates and purified linear tetra-ubiquitin chains *in vitro*. Additionally, we provide evidence that caf1 binds to 19S regulatory particle complex of 26S proteasome.

Taken together we conclude that Caf1 is a novel ubiquitin binding protein, functions as an ubiquitin shuttle factor required for degradation of ubiquitin-modified proteins at the post ubiquitylation step.

## Materials and Methods

### Yeast media

Standard growth medium including YP and synthetic dropout (SD) having either 2% glucose or galactose with specific amino acid(s) and nucleotide(s) drop outs were prepared and used in this entire study [36].

### Strains and plasmids

In the current study BY4741 wild type yeast strain and its mutant derivatives were used. The genotypes of all the strains are provided in the strain table 1 attached in the supplemental information section. We generated a stable *TRP1* gene deletion in BY4741 wild type yeast strain using a previously described method [37]. To introduce *TRP1* gene deletion chromosomally the pNKY1009 plasmid was digested with BglII and EcoRI to release the deletion cassette. The released fragment carrying the URA3 marker with flanking TRP1 gene sequences were transformed into the respective yeast strains and Ura^+^ and Trp^−^ colonies were selected on agar plates containing appropriate medium. In the next step, the *URA3* gene was cured out by selection on SD plates containing 5-Fluorooroticacid hydrate(1g/l, Sigma) as described previously. The *caf1* gene was deleted in PYGA12 strain using S1 and S2 primers as described earlier [38].

**Table1:** Yeast strains

| Yeast strain | Genotype | Source |
| --- | --- | --- |
| BY4741-WT | *MATa his3-Δ1 leu2Δ0 met15-Δ0 ura3-Δ0* | Euroscarf |
| *caf1Δ* | *MATa his3-Δ1 leu2Δ0 met15-Δ0 ura3-Δ0 caf1::KanMX4* | Euroscarf |
| *not1-ts* | *MATa his3-Δ1 leu2Δ0 met15-Δ0 ura3-Δ0 not1-ts::KanMX4* | Euroscarf |
| *not2-ts* | *MATa his3-Δ1 leu2Δ0 met15-Δ0 ura3-Δ0 not2-ts::KanMX4* | Euroscarf |
| *not3Δ* | *MATa his3-Δ1 leu2Δ0 met15-Δ0 ura3-Δ0 not3::KanMX4* | Euroscarf |
| *not5Δ* | *MATa his3-Δ1 leu2Δ0 met15-Δ0 ura3-Δ0 not5::KanMX4* | Euroscarf |
| *caf40Δ* | *MATa his3-Δ1 leu2Δ0 met15-Δ0 ura3-Δ0 caf40::KanMX4* | Euroscarf |
| *caf130Δ* | *MATa his3-Δ1 leu2Δ0 met15-Δ0 ura3-Δ0 caf130::KanMX4* | Euroscarf |
| *ump1Δ* | *MATa his3-Δ1 leu2Δ0 met15-Δ0 ura3-Δ0 ump1::KanMX4* | Euroscarf |
| PYGA12 | *MATa his3-Δ1 leu2Δ0 met15-Δ0 ura3-Δ0 trp1Δ* | This study |
| YAP16 | *MATa his3-Δ1 leu2Δ0 met15-Δ0 ura3-Δ0 trp1Δ KanMX4::P<sub>GAL1</sub>-PRE2* | This study |
| YAP19 | *MATa his3-Δ1 leu2Δ0 met15-Δ0 ura3-Δ0 trp1Δ caf1::His3MX6* | This study |
| *ufd4Δ* | *MATa his3-Δ1 leu2Δ0 met15-Δ0 ura3-Δ0 ufd4::KanMX4* | Euroscarf |

**Table 2A:**
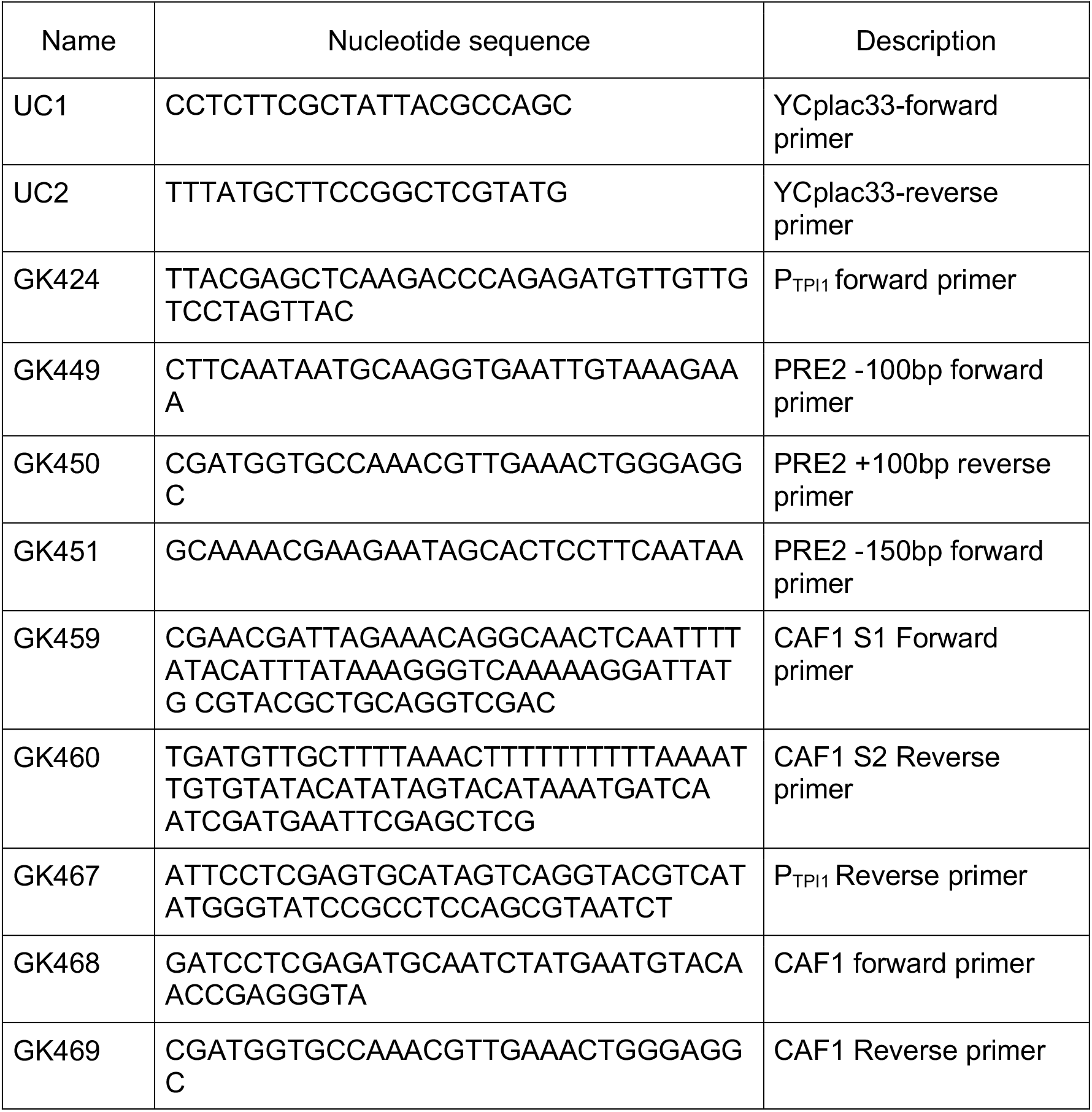
Primers

**Table 2B:** Plasmids

| Plasmid Name | Description | Reference/<br>Sources |
| --- | --- | --- |
| Ycplac33 | CEN/ARS, URA3, AmpR | [49] |
| Ycplac22 | CEN/ARS, TRP, AmpR | [49] |
| Ycplac111 | CEN/ARS, LEU, AmpR | [49] |
| YEP105 | P<sub>CUP1</sub>- MYC-UB, 2μ, TRP, AmpR | [41] |
| pMAF18 | P<sub>CUP1</sub>- Ub-V<sub>76</sub>-eKa-HA-URA3, CEN/ARS, LEU, AmpR | [13] |
| pMAF17 | P<sub>CUP1</sub>- Ub-R-eKa-HA-URA3, CEN/ARS, LEU, AmpR | R. J. Dohmen |
| pD5258 | P<sub>ODC</sub>-ODC1-42-URA3-2xHA, CEN/ARS, LEU, AmpR | [40] |
| pMAF56 | P<sub>CUP1</sub>-DHFRWT-2HA, CEN/ARS, LEU, AmpR | R. J. Dohmen |
| pMAF56C | P<sub>CUP1</sub>-DHFR<sup>mutC</sup>-2HA, CEN/ARS, LEU, AmpR | R. J. Dohmen |
| pAK29 | P<sub>TPI1</sub>-2HA-CAF1- CEN/ARS, LEU, AmpR | Current study |
| pBA2VV | P<sub>CUP1</sub>- Ub-V<sub>76</sub>-eKa-HA-URA3, TRP/ARS, LEU, AmpR | R. J. Dohmen |

By using S1 and S4 primers (GK449 and GK450), which carries PRE2 gene flanking regions, the KanMX4::P_*GAL1*_-PRE2 cassette was amplified by PCR using genomic DNA isolated from YGA140 strain. PCR product was used to transform PYGA12 strain and after overnight recovery in YPGAL medium, the transformants were selected on YPGal-G418 plates (300μg/ml).The resulting YAP16 strain was verified by absence of growth in YP-glucose plates. Plasmids pMAF17 (P_*CUP1*_-UB-R-eKa-HA-URA3), pMAF18 (P_*CUP1*_-UB-V^**76**^-eKa-HA-URA3), pMAF56 (P_*CUP1*_-DHFR^WT^-2HA) and pMAF56C (P_*CUP1*_-DHFR^mutC^-2HA), were derived from the *CEN/LEU2* vector pRS315, and the plasmid pBA2VV, is a derivative of *CEN/TRP1* vector pRS314 were also kind gifts from Dohmen RJ. pDG258 (P_*SPE1*_-ODS-URA3-2HA) is derived from *CEN/LEU2* vector pPM91 as described [39]. pAK29 (P_*TPI1*_-2HA-CAF1-T_*CYC1*_) is derived from PAK28 which is constructed on *CEN/LEU2* vector YCplac111. For constructing pAK29 the vector backbone was obtained by double digesting PAK28 clone with SacI and XbaI restriction enzymes. Then the insert one carrying P_*TPI1*_-2HA was amplified from pAK28 plasmid template using GK424 and GK467 primer pairs and double digested with SacI and XhoI. Insert two carrying CAF1 gene fragment was generated from genomic DNA template isolated from BY4741 strain using GK468 and GK469 primer pairs and double digested with XhoI and XbaI enzymes. The digested vector and inserts were subjected to three part ligation using T4 DNA ligase. YEp105 (expressing MYC-UB from P_*CUP1*_) is a *2μ/TRP1* plasmid [40]. Plasmid pGA54 (expressing recombinant linear tetra-ubiquitin in *Escherichia coli*) was a kind gift from Dohmen RJ. The plasmids and primers used in this study are listed in the supplementary table 2 A and B.

### Yeast growth condition

Wild type and mutant yeast strains were cultured at 25°C in the YP/SD agar plates or in liquid medium containing either glucose or galactose as a carbon source. For essential gene mutants, yeast strains carrying the temperature sensitive mutant alleles were grown at non-permissive temperature (37°C). The specific growth conditions are described in the respective experiments in the results section.

### Western blot analysis for protein detection and quantification

The steady state levels of various substrate proteins were analyzed on western blot. The *S. cerevisiae* transformants were inoculated and grown overnight (ON) at 25°C in SD medium lacking respective amino acids like Leucine (SD –Leu) as per the requirements. Next day, the ON cultures were diluted in fresh SD –Leu medium (OD_600_ - 0.2) having 100μM CuSO4 as mentioned. Cell cultures were grown until mid-log phase (OD_600_ - 0.8 to 1.0) then the cells corresponding to 5.0 OD_600_ were harvested. Total cell extracts were prepared as described earlier [41], the cell pellets were washed once with water, then mixed with 250 μl of ice cold 1.85N NaOH. After incubation for 10 min on ice 250 μl of 50% TCA was added to each sample and mixed well then cells were pelleted by centrifuging at 14,000 rpm for 10 min. The supernatant was discarded and the cell pellets were re-suspended in 100 μl of 1M Tris (pH 7.5) and pelleted again as mentioned above. Finally, the pellets were re-suspended in 1x Laemmli loading buffer (0.0625M Tris–HCl (pH 6.8), 2% SDS, 1% β-mercaptoethanol, 10% glycerol, 0.002% bromophenol blue) and lysed by boiling at 85°C for 5 min. After a brief centrifugation to pre-clear the cell debris, the yeast cell extracts corresponding to 0.416 OD_600_ were loaded onto a SDS-PAGE and western blotting was done to detect specific proteins. Quantification of the corresponding signals were performed as described earlier [42]. Anti-Ha (C29F4 from cell signalling technologies, 1:2000 dilution) was used for detecting ha-tagged proteins. Anti-Rpt5 (Enzolife sciences, 1:1000 dilution) was used to detect the 19S regulatory particle subunit of yeast 26S proteasome. Anti-Tpi1 (Lab source, 1:10000 dilution) and Anti-Pgk1 (Novex, 1:5000 dilution) were used to detect loading control proteins Tpi1 and Pgk1. Anti-mouse or anti-rabbit IgG coupled to peroxidase were used as secondary antibodies (Millipore, 1:5000 dilution). Western bright ECL substrate or western bright Sirius-femtogram HRP (Advansta) was used for developing the blots. Specific signals were captured using a Bio-Rad ChemiDoc™ MP Imaging system and quantified using the software Bio-Rad Image Lab 5.2.1.

### Cycloheximide chase analysis to monitor protein turn-over rate

Proteins half-life were determined by inhibiting *de novo* protein translation. Exponentially growing yeast cells were treated with protein translation inhibitor cycloheximide at a concentration of 100 mg/l, samples were collected at the indicated time points. Then samples were processed and analyzed on SDS-PAGE followed by western blotting as described above. Quantification of western blots signals were performed as described earlier [42].

### Immunoprecipitation

For Co-Immunoprecipitation (Co-IP) experiments the plasmids YEp105 and pAK29 expressing MYC-UB and/or 2HA-CAF1, or its isogenic empty vector plasmids pPM91, YEplac112 were used for transforming YAP16 yeast strain (BY4741,KanMX4:: P_*GAL1*_-PRE2). Transformants were selected on SGAL-Leu-Trp agar plates. Primary cultures were grown in galactose containing SGAL-Leu-Trp medium. For proteasomal inactivation the PRE2 expression from P_*GAL1*_ was turned off by diluting the primary cultures into a fresh liquid medium containing 2% glucose and grown at 25°C for 12 hrs. For Co-IP, the cells corresponding to 80 OD_600_ were harvested and the pellets were frozen at −65°C until further usage. For lysis at native condition pellets were re-suspended in 600 μl of lysis buffer (50mM HEPES pH7.5, 150mM NaCl, 5mM MgCl_2_, 1% Triton X-100, 40mM N-Ethylmaleimide, EDTA-free protease inhibitor cocktail (Roche) and 2mM PMSF (Sigma/Merck). Afterwards 300 μl of acid washed glass beads (425-600 μM from Sigma/Merck) were added to the cell suspension and kept in ice for 5 min then tubes were vortexed three times with 1 min incubation on ice between each round. Cell debris and glass beads were pelleted down by centrifugation at 14,000 rpm for 10 min at 4°C. Equal amounts (Protein estimated by Bradford) of soluble fractions were incubated with 40 μl of pre-equilibrated anti-HA affinity agarose gel (Roche) in binding buffer (Lysis buffer). Binding was done at 4°C for 3hrs then beads were washed 3X 5 min with binding buffer with constant changing of Eppendorf tubes in-between to reduce the non-specific binding. Bound proteins were eluted by boiling at 85°C for 5 minutes with 1X LLB. Eluates were analyzed by SDS-PAGE followed by western blotting. Anti-Ha (Cell

Signaling Technologies,1:5000 dilution) or anti-Myc (9B11, Cell Signaling Technologies, 1:1000 dilution) or anti-Rpt5 (Enzolifesciences, 1:1000) was used for protein detection. To perform denaturation immunoprecipitation of Ha2-Caf1 soluble cell lysates were prepared as described above but SDS (2%) as well as β-mercaptoethanol (1%) was added. The samples were then boiled at 95°C for 5min. Next, the samples were diluted to reduce the SDS and β - mercaptoethanol to 0.05% and 0.025% respectively. Diluted samples were incubated with 60 μl of pre-equilibrated anti-Ha affinity agarose gel and washed prior to elution as described above.

### Ubiquitin binding assay

The YAP16 yeast strain was transformed either with PAK29 plasmid or with its isogenic empty vector. Transformants were selected on SGAL-Leu agar plates and the expression of 2HA-CAF1 from PAK29 plasmid was induced with 2% galactose. Simultaneously the *E. coli* BL21 codon plus strain expressing the recombinant 6-His linear tetra ubiquitin was purified using nickel NTA resin. After induction with 1mM IPTG, 1 l *E.coli* culture was harvested and re-suspended in the lysis buffer (50mM HEPES pH7.5, 150mM NaCl, 5mM MgCl_2_, 1% Triton X-100, EDTA-free protease inhibitor cocktail (Roche) and 2mM PMSF (Sigma/Merck). Next, soluble cell extracts were prepared by glass bead lysis method and the soluble *E. coli* extract containing the linear tetra ubiquitin was bound to 6ml of preequilibrated nickel NTA beads at 4°C for 4 hrs. After binding step the beads were extensively washed with lysis buffer supplemented with 10mM imidazole to reduce the non-specific binding. Finally, the nickel NTA beads bound linear tetra ubiquitin was eluted with 500mM imidazole with PH 7.5 for 1 hour at 4°C. For Co-IP experiment the YAP16 yeast transformants corresponding to 100 OD_600_ was harvested and stored at −65°C until further use. Immunoprecipitation of Ha2-Caf1 was done as described in the earlier section. After extensive washing steps, the resin bound Ha tagged Caf1 was incubated with recombinant linear tetra ubiquitin purified from *E.coli*, the binding was done at 4°C for 1 hour. Later, the beads were extensively washed and the bead bound Ha tagged Caf1 protein was eluted by boiling with 1x LLB at 85°C for 5 min.

## Results

### Ubiquitin-dependent substrate degradation by the proteasome is impaired upon *caf1* deletion

CCR4-NOT is a multi-subunit protein complex that functions in several regulatory pathways [16]. One of the interesting observations from the earlier studies is that CCR4-NOT complex subunits interact with 26S proteasome and molecular chaperones [13,31,32], however, the functional relevance of such interactions are largely unexplored. These observations prompted us to postulate CCR4-NOT subunits might have function in the protein degradation pathway. In order to identify the relevant CCR4-NOT complex proteins we analyzed the steady state levels of well-established N-end Rule substrate (R-Ura3) and UFD substrate (Ub^V76^-Ura3) of UPS (Fig 1). As a control we took wild type (WT) and *ump1* deletion strains (Ump1 protein is required for assembly and maturation of proteasome [43]. Our experiment surprisingly reveled that *caf1* deletion results in increased accumulation of UFD substrate Ub^V76^-Ura3 and R-Ura3 substrates (Figure 2A & B) as compared to the wild type strain, however, as expected *ump1* deletion strain showed the stronger accumulation of both substrates due to impaired proteasomal function (Figure 2A & B). Next, we systematically tested the stable deletion mutants (*not3, not5, caf40* and *caf130*) as well as the conditional mutants (*not1-ts and not2-ts*) for their requirement in the protein degradation pathway and we found that most of the CCR4-NOT complex subunit mutants are dispensable for the proteolysis of tested ubiquitin-dependent proteasomal substrates (Figure 3A). Next, we made use of the Ura3 reporter protein that is present in the UFD substrate and further tested the requirement of Caf1 protein in the Ub^V76^-Ura3 substrate proteolysis by performing growth assay experiment in WT and *caf1Δ* mutant strain, as a control we took *ump1Δ* and *ufd4Δ* mutant strains (UFD4 is the E3 ligase for UFD substrate). The principle behind this experiment is that WT and all the mutant strains are Uracil auxotrophs, only the mutant strains that is defective in degrading Ub^V76^-Ura3 substrate would produce enough Ura3 reporter protein in the cell and supports cell viability in medium lacking Uracil. As reported earlier, the *ump1Δ* and *ufd4Δ* mutant strains accumulated Ub^V76^-Ura3 substrate and showed rapid growth [44], similarly, consistent with our steady state experiment result, *caf1* deletion accumulated Ub^V76^-Ura3 substrate and significantly suppressed the uracil auxotrophy phenotype (Figure 3B). In addition, the ectopic expression of wild-type CAF1 complemented the proteolysis defects of *caf1Δ* mutant strains (Figure 2E). We considered two possible explanation for the increased steady state levels of Ub^V76^-Ura3 and R-Ura3 substrate in *caf1Δ* strain, either Caf1 has a direct role in the protein degradation pathway, thus deletion of *caf1* resulted in defective proteolysis and accumulation of Ub^V76^-Ura3 and R-Ura3 substrates or its deletion phenotype caused higher translation rates of Ub^V76^-Ura3 and R-Ura3 proteins as well as higher transcription from *CUP1* promoter. In order to test whether *caf1Δ* strain changes *CUP1* promoter expression levels, we tested the DHFR reporter protein levels driven by same promoter in WT, *caf1Δ* and *ump1Δ* strains and found that the expression of wild-type DHFR was similar in all the tested strains (Supplementary figure 2), indicating that *caf1Δ* has no effect on *CUP1* promoter expression levels. Next, we explored our analysis with additional proteasomal substrates such as misfolded Dhfr^mutC^ protein which is degraded by UPS mediated protein quality control pathway and ODS-Ura3 protein (ODS-ODC degradation signal), degraded by ubiquitin-independent pathway [39]. Our steady state analysis showed that *caf1Δ* strain did not accumulate ODS-Ura3 protein and Dhfr^mutC^ (Misfolded protein) (Figure 2C and D). Further, we confirmed the Caf1 proteolytic function by monitoring the degradation kinetics of Ub^V76^-Ura3, Dhfr^mutC^ and ODS-Ura3 proteasomal substrates using protein translation inhibitors. Our cycloheximide chase assay revealed that Ub^V76^-Ura3 substrate was stabilized upon *caf1* deletion (Figure 4), however, the proteolysis of ODS-Ura3 and Dhfr^mutC^ substrates were normal and similar to wild type strain (supplementary figure 1A and B). *ump1* deletion showed stabilization of all tested proteasomal substrates (Figure 4 and supplementary figure 1 A and B), indicating the requirement of functional 26S proteasome for substrate proteolysis. Altogether, these results implicate that Caf1 plays a novel role in the UPS pathway specifically required for the proteolysis of selective ubiquitin-dependent substrates, but it is dispensable for the degradation of ubiquitin-independent substrates.

**Figure 1:**
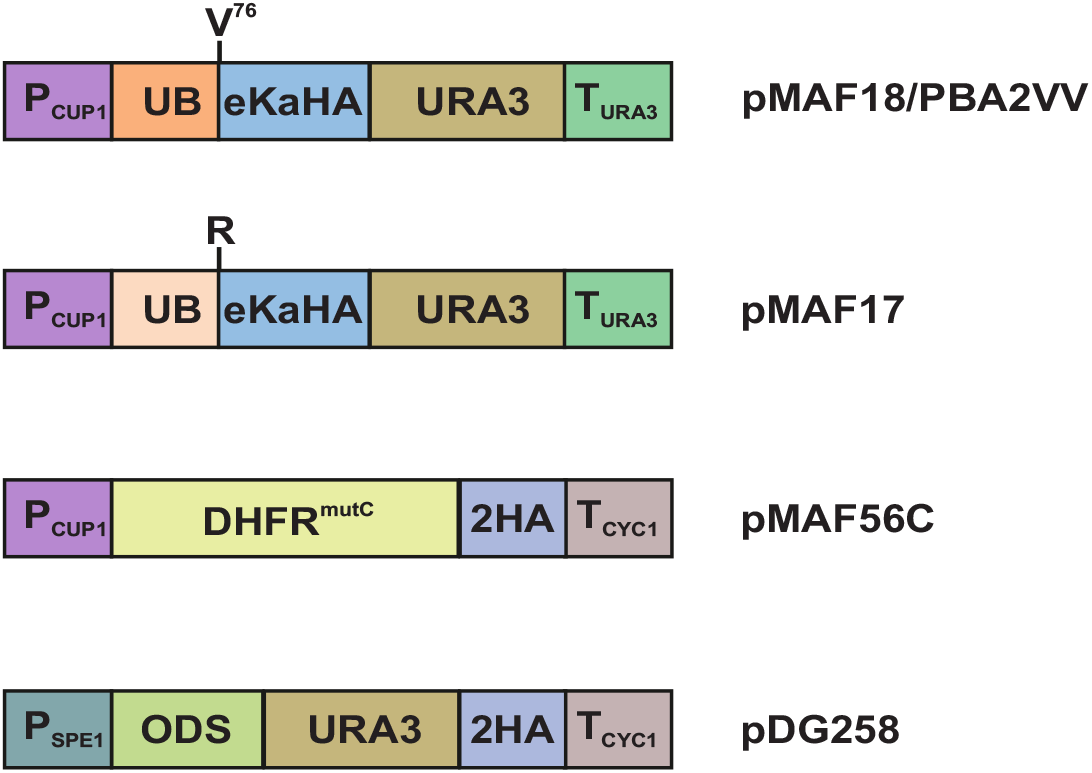
Graphical representation of different model substrates of UPS pathway. Shown is the detailed illustration of proteasomal substrate carrying specific proteolytic signals. The construct pMAF17 and pMAF18/pBA2VV producing R-Ura3 and Ub-V^**76**^-Ura3 proteins are ubiquitin-dependent substrates degraded by N-end rule and Ubiquitin Fusion Degradation (UFD) pathways respectively. The folding incompetent mouse Dhfr^mutC^ (pMAF56C) is a substrate of protein quality control pathway. ODS-Ura3 protein carrying the N-terminal unstructured region of yeast ornithine decarboxylase (ODC) is an ubiquitin-independent pathway substrate of the proteasome.

**Figure 2:**
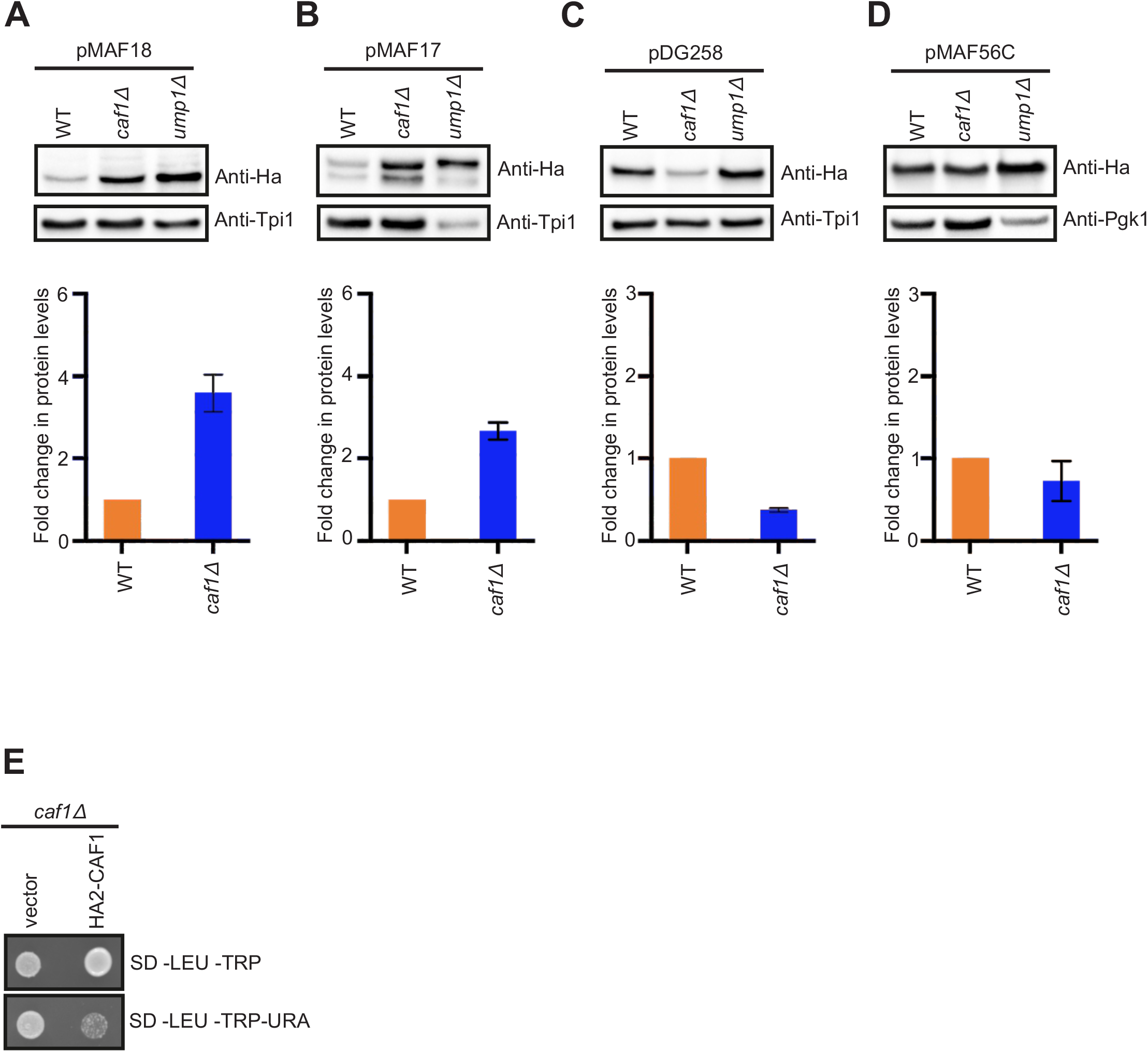
Steady state levels of different proteasomal substrates in *caf1D* strain. Western blot signals showing the steady state protein levels of (A) Ub-V^**76**^-Ura3, (B) R-Ura3, (C) ODS-Ura3 and (D) misfolded Dhfr^mutC^ substrates in WT, *caf1Δ* and *ump1Δ* strains (upper panel). Anti-Ha antibody was used to detect the protein levels of various proteasomal substrates. Pgk1 or Tpi1 were used as protein loading controls (upper panel). Lower panel shows densitometric analysis of protein signals in WT and *caf1Δ* strains. Error bars denote s.d; n≥3. (E) Shown is the Ura3 based reporter protein complementation assay in *caf1Δ*. Cells corresponding to 10.0/ml OD_600_ of 3μl cultures was spotted on selection plates as indicated and growth patterns are documented after 3-4 days of incubation at 25°C. Represented data is the sample volume of four different biological repeats.

**Figure 3:**
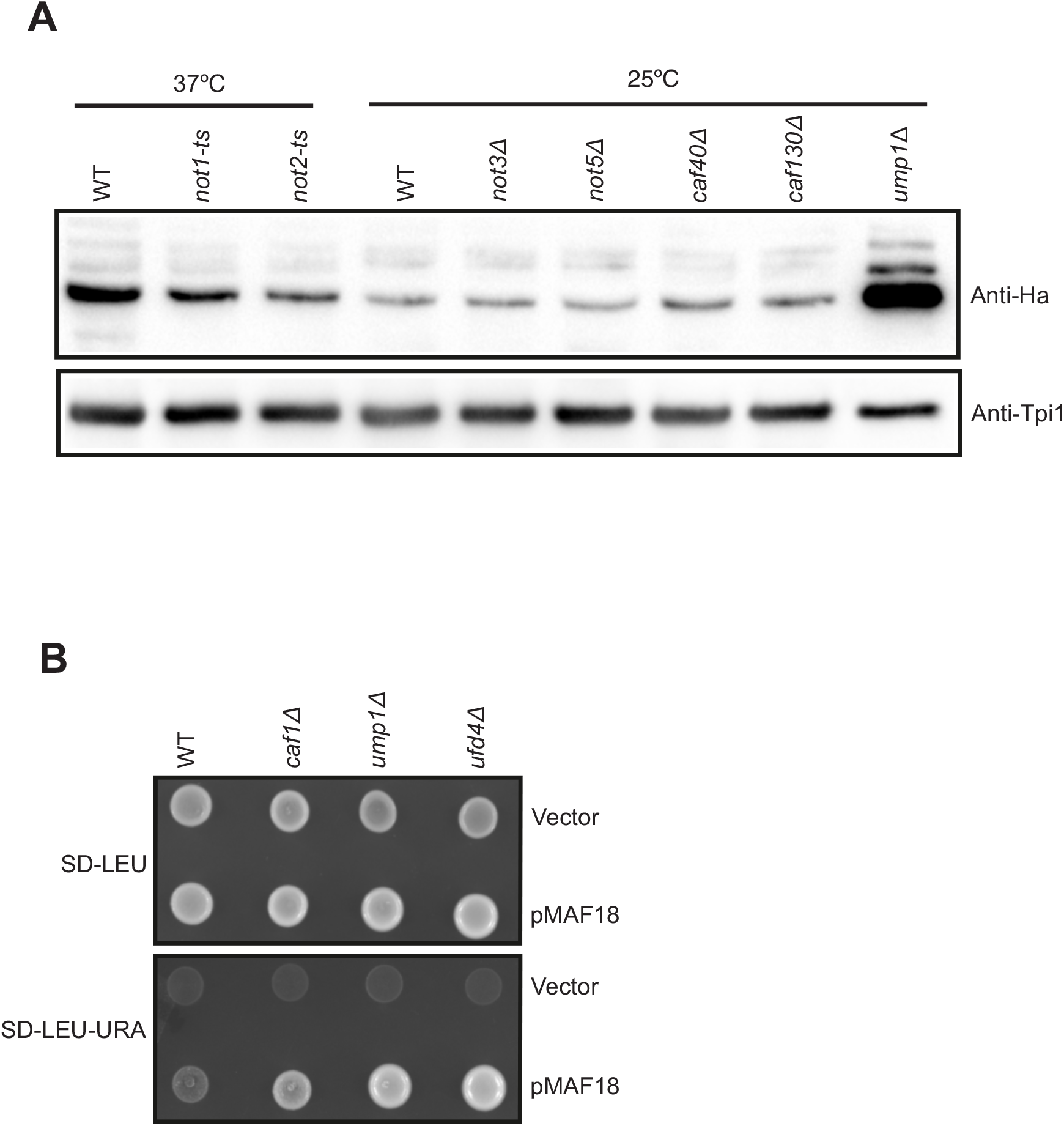
Proteolysis of Ub-V^76^-Ura3 substrates in different CCR4-NOT complex subunit mutants. (A) Data displayed is the steady state Ub-V^76^-Ura3 protein levels in WT, *not1-ts, not2-ts, not3Δ, not5Δ, caf40Δ, caf130Δ, and ump1Δ*, mutant strains. All the strains are grown at 25°C, except, the temperature sensitive mutants *not1-ts* and *not2-ts* were transiently shifted to 37°C for 1hr to inactivate Not1 and Not2 protein function. Anti-Ha and anti-Pgk1 antibodies were used for immuno blotting. Data shown is the representation of three independent biological samples. (B) Growth assay pattern of WT, *caf1Δ, ump1Δ*, and *ufd4Δ* strains expressing UFD pathway substrate (Ub-V^76^-Ura3, pMAF18). Uracil auxotrophy phenotype of the above indicated yeast transformants were documented after 3-4 days of growth at 25°C.

**Figure 4:**
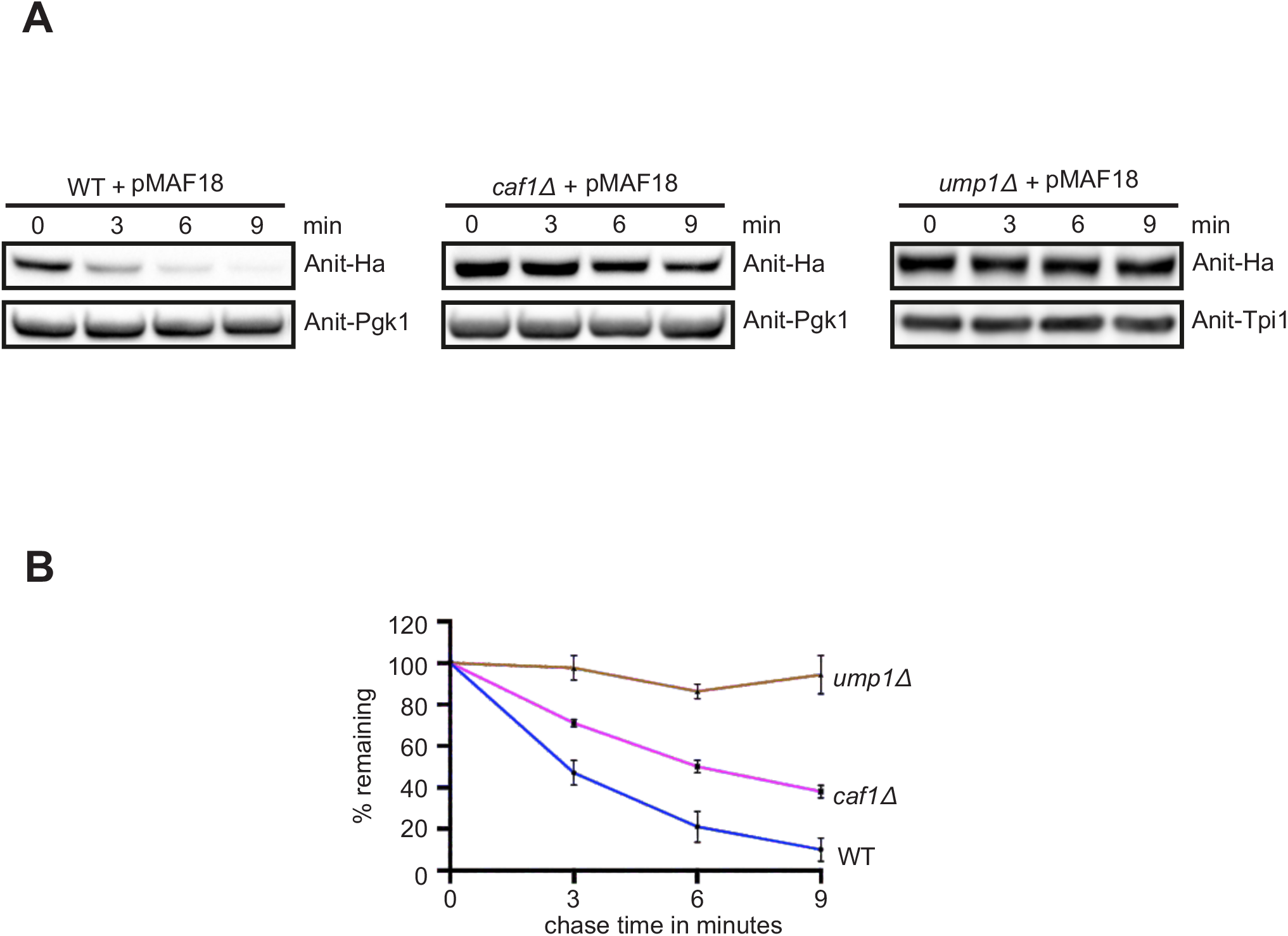
Degradation analysis of UFD substrate in *caf1Δ*. (A) Logarithmically grown yeast cells were treated with protein translation inhibitor cycloheximide to monitor the degradation kinetics of Ub-V^76^-Ura3 protein in WT, *caf1Δ* and *ump1Δ* strains. Cells were harvested at specific time points as indicated and total cell extracts were analyzed by western blotting method. Anti-Ha, anti-Tpi1, and anti-Pgk1 antibodies were used in the western blot analysis. (B) Quantification of Ub-V^76^-Ura3 protein signals in WT *caf1Δ*, and *ump1Δ* strains. Error bars denote s.d; n?3

### Caf1 accumulates ubiquitin-modified UFD substrate and interacts with 19S cap of 26S proteasome

After finding that Caf1 protein is involved in the UPS pathway we postulated Caf1 might function either at pre and/or post ubiquitylation steps in the UFD substrate degradation. In order to test our idea we analyzed the ubiquitylation pattern of UFD substrates (UFD substrate was shown to accumulate ubiquitin-modified form when proteasome activity is impaired [43]. Our results showed that in *caf1Δ* strain UFD substrate accumulates as ubiquitin-modified form similar to the proteasome defective *ump1* mutant in lower percentage SDS PAGE gels followed by western blotting (Figure 5A). From this result we concluded that Caf1 functions at the post ubiquitylation step to degrade UFD substrate. Intrigued by the above result we then tested if Caf1 would physically interact with proteasome and more specifically with 19S cap subunits of the 26S proteasome. In order to address that we have immunoprecipitated Caf1 and intriguingly found that 19S RP cap subunit Rpt5 co-purified with Caf1 (Figure 5B) suggesting that Caf1 indeed interacts with proteasome and thereby promotes the degradation of ubiquitin-modified proteins in the UPS pathway.

**Figure 5:**
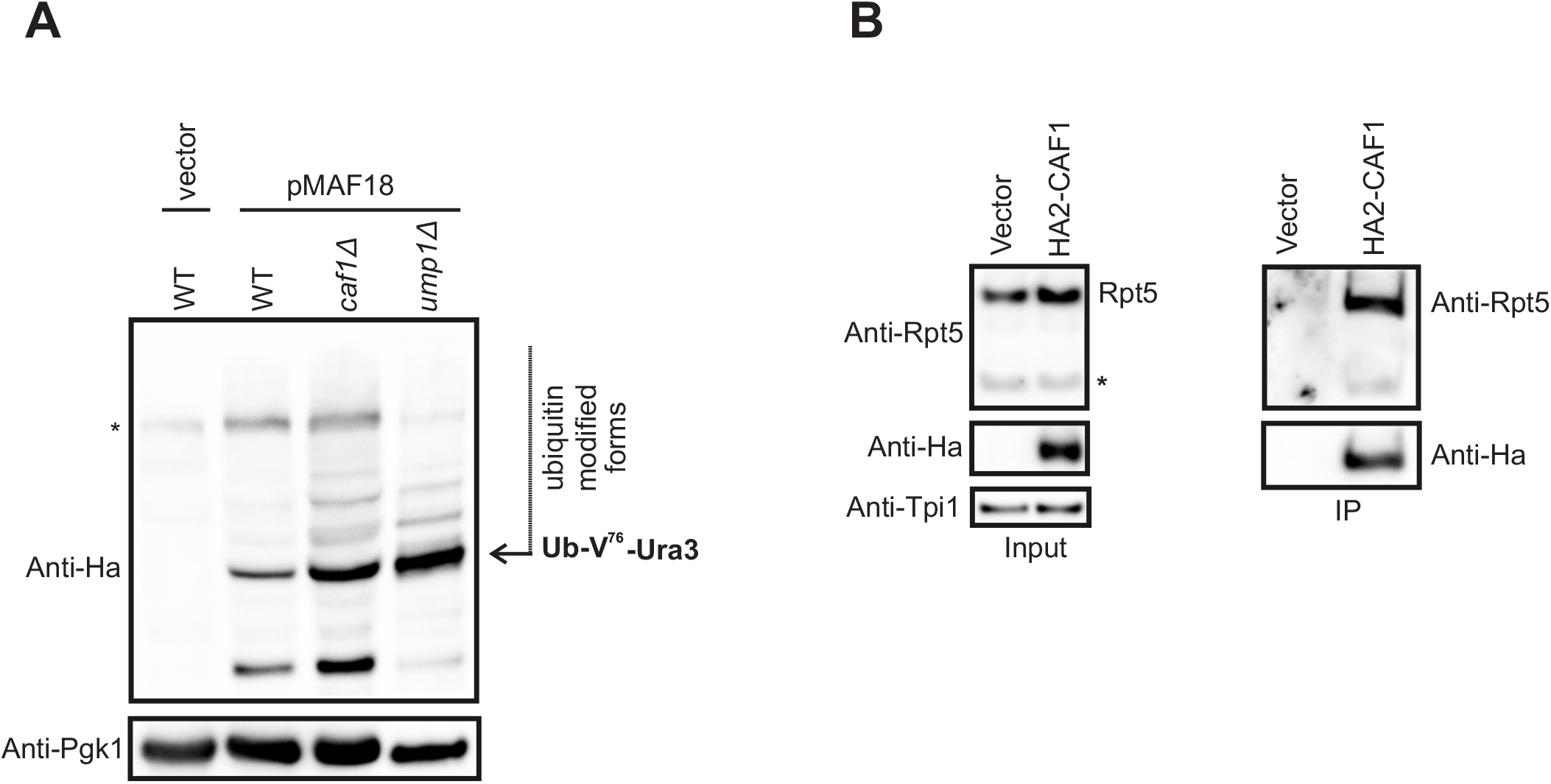
Caf1 functions at post-ubiquitylation step and associates with 19S regulatory particle complex. (A) Immuno blot analysis showing poly-ubiquitylated species of UFD (Ub-V^76^-Ura3) substrate in WT, *caf1Δ*, and *ump1Δ* strains. Data represents three independent biological repeats. Anti-Ha was used to detect Ub-V^76^-Ura3 substrate. Pgk1 was used as a loading control. Asterisk denotes cross-reactive bands. (B) Co-immuno precipitation experiment showing interaction of Ha2-Caf1 with 19S regulatory particle complex. YAP16 carrying empty plasmid vector was used as a control. Immunoblot analysis of input (left panel) and IP material (right panel). Anti-Tpi1 antibodies was used as a protein loading control. AntiHa and anti-Rpt5 antibodies were used to decorate Ha2-Caf1 and Rpt5 proteins respectively. Asterisk denotes degradation product of Rpt5 protein. Data shown is the representation of three independent experiments.

### Caf1 binds to ubiquitin conjugates and interacts with linear tetra ubiquitin

Degradation of poly-ubiquitylated proteins by 26S proteasome initially requires the docking of substrate proteins with 19S regulatory particle complex of 26 proteasome, canonical ubiquitin shuttle factors on the one hand bind to proteasome and on the other hand interacts with ubiquitin chains of the substrate proteins thereby functioning as bridging factor between the poly-ubiquitylated proteins and 26S proteasome [10,11]. In line with that, we hypothesized Caf1 might bind to ubiquitin conjugates in order to function as a shuttle factor protein. The above hypothesis was validated by immunoprecipitation experiment, we intriguingly found that cellular ubiquitin conjugates co-purified along with Caf1 protein under native condition (Figure 6A). The above finding raises a concern Caf1 might itself get ubiquitylated and contributed to the observed ubiquitin conjugates signal in our CO-IP experiment. To test, if Caf1 is ubiquitylated, we performed immuno-precipitation experiment under denaturation conditions and found no significant ubiquitylation of Caf1 protein (Figure 7). Following this exciting observation we had asked if Caf1 protein would be able to interact with linear tetra ubiquitin chains. Surprisingly, from our partial *in vivo* and *in vitro* experiment data we found that indeed Caf1 interacts with recombinant linear tetra ubiquitin chains purified from *E.coli* (Figure 6B). From these findings we conclude that Caf1 is a novel ubiquitin binding protein, it binds cellular ubiquitin conjugates and therefore shuttles it to 26S proteasome for proteolysis.

**Figure 6:**
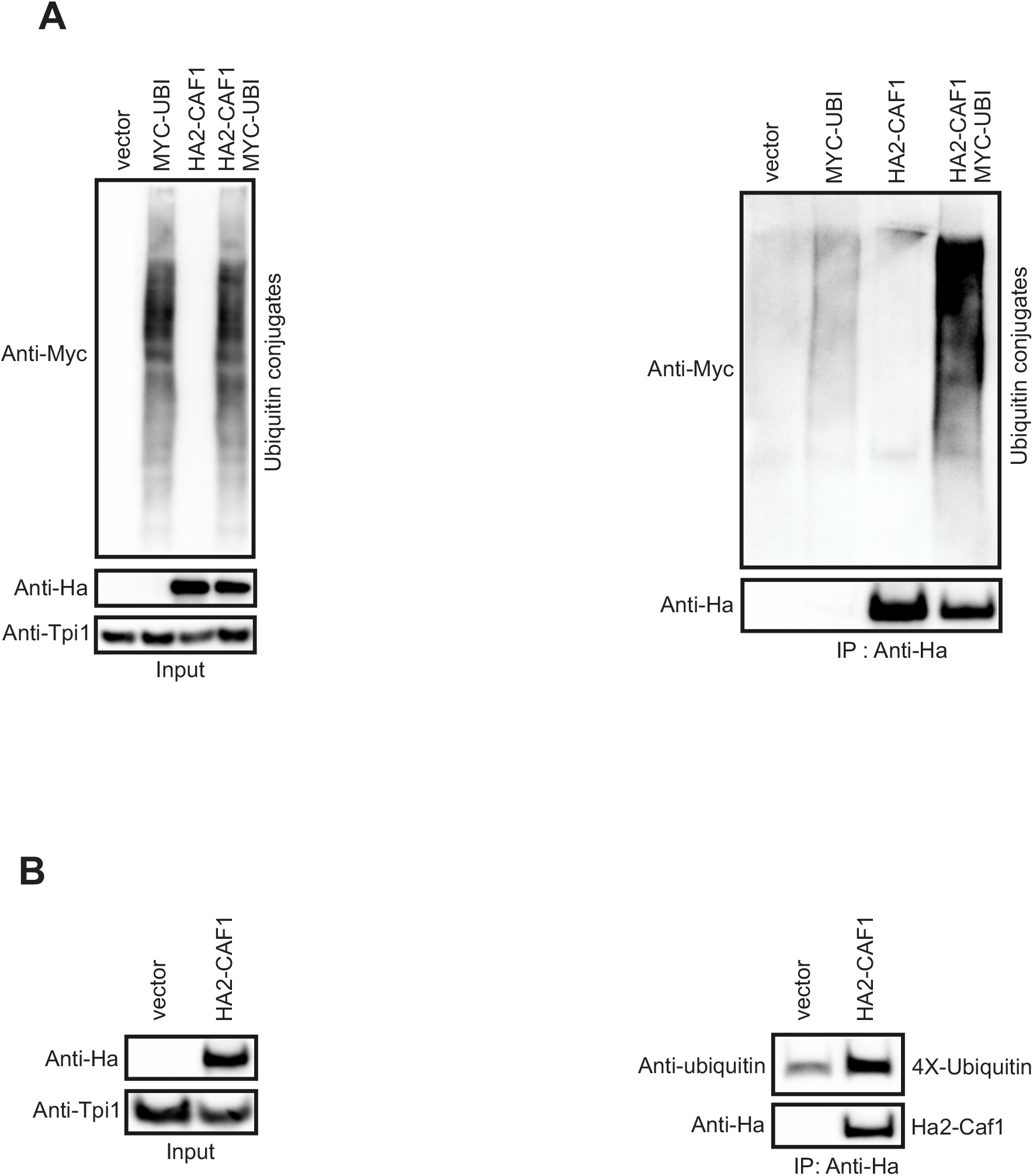
Caf1 interacts with cellular ubiquitin conjugates and linear tetra ubiquitin. (A) Western blot showing association of high molecular weight ubiquitin conjugates with Ha2-Caf1 protein under native conditions. YAP16 strain transformed with empty plasmid vector was used as a control. Immuno blot analysis showing input material (left panel) and IP material (right panel). Anti-Myc was used to detect ubiquitin conjugates. Anti-Ha and anti-Tpi1 antibodies were used to decorate Ha2-Caf1 and Tpi1 proteins respectively. Data represents results from three independent experiments. (B). Partial *in vivo* and *in vitro* experiment showing the interaction of Ha2-Caf1 protein with linear tetra ubiquitin chains purified from *E.coli*. Shown are the western blot analysis of input (left panel) and IP materials (right panel). Anti-ubiquitin antibody was used to detect linear tetra ubiquitin chains. Caf1 and Tpi1 protein levels were detected using anti-Ha and anti-Tpi1 antibodies respectively. Data shown is the representation of results from three independent experiments.

**Figure 7:**
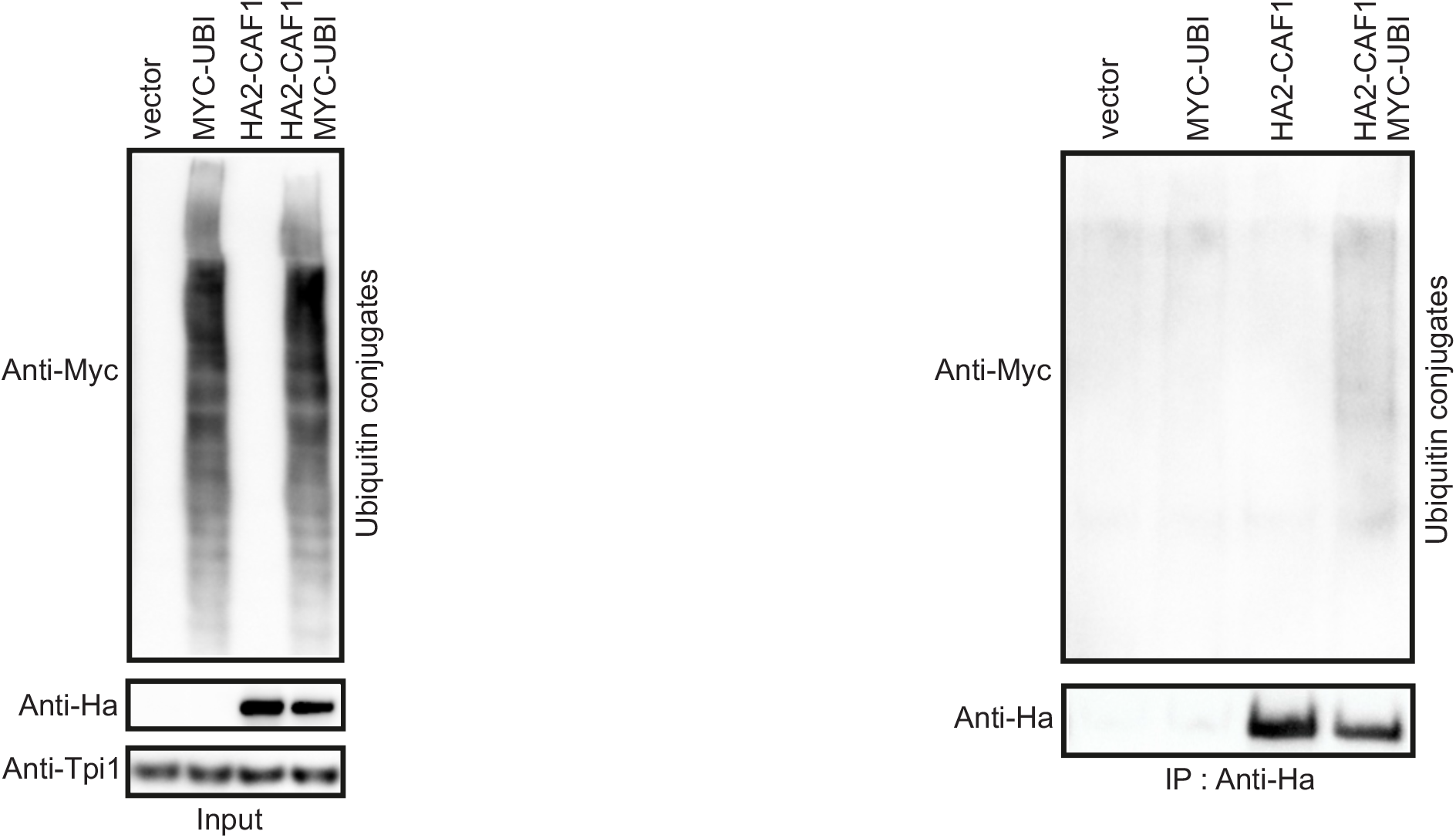
Denaturing immunoprecipitation of Ha2-Caf1 protein. Western blot showing immunoprecipitation of Ha2-Caf1 protein under non-native conditions. Denaturation IP was performed as detailed in methods section. The eluted IP material was loaded on SDS gel and followed by immuno detection using anti-Myc and anti-Ha antibodies. Left panel shows input levels and right panel shows eluted material. Tpi1 was used as a protein loading control. Data represents repetition of three independent experiments.

## Discussion

CCR4-NOT is a highly conserved multisubunit protein complex plays a vital role in maintaining RNA homeostasis in the cells [15,16]. The role of CCR4-NOT complex also extends beyond mRNA quality control pathway, in yeast the CCR4-NOT complex subunits Not4, Ccr4 and Not1 were shown to interact with 26S proteasome and molecular chaperones [13,31,32,45], these findings supports that CCR4-NOT complex might function in the protein degradation pathways. Therefore, in the current study we have methodically tested the functional requirement of different CCR4-NOT complex subunits Caf1, Caf130, Caf40, Not1, Not2, Not3 and Not5 in the UPS pathway. Based on our evidence we report that the Ccr4 associated factor Caf1 has a previously unidentified role in the UPS pathway. We found that *caf1* deletion results in accumulation of UFD pathway substrates (UbV^76^-Ura3) (Fig 2A), in support of that *caf1D* mutant strains suppressed the uracil auxotrophy phenotype due to the higher levels of intracellular Ura3 reporter proteins (Figure 2E and 3B). These findings arises a concern that the increased levels of UbV^76^-Ura3 proteins in *caf1D* might be the consequence of increased CUP1 promoter driven expression of UFD substrate, however, straight in contrast our control experiment proved that the CUP promoter driven expression of stable Dhfr protein levels were similar in WT and *caf1D* mutant strain (Supplementary figure2). Importantly, our cycloheximide chase analysis experiment revealed that Caf1 protein indeed has a direct role in the UFD pathway substrate proteolysis (Figure 4). Additional support on the proteolytic role of Caf1 protein in the UPS pathway was demonstrated by the accumulation of N-end rule pathway substrate R-Ura3 in *caf1D* mutant strain (Figure 2B). Strikingly, we observed that *caf1* deletion accumulated Ub-V^76^-Ura3 substrates in the poly-ubiquitylated forms (Figure 5A). Based on the above discovery we conclude that Caf1 functions at the post-ubiquitylation step in targeting ubiquitin-modified Ub-V^76^-Ura3 substrate proteins for proteasomal degradation. Our extended degradation analysis of multiple proteasomal substrates revealed that Caf1 is dispensable for the proteolysis of ubiquitin-independent ODS-Ura3 and ubiquitin-dependent folding deficient Dhfr^mutC^ proteasomal substrates (supplementary Figure 1 A and B). These results indicate that *caf1D* does not affect the UPS function largely, hence, the degradation of several tested proteasomal substrates are largely unaffected in our experimental conditions. We speculated that the potential post-ubiquitylation role of Caf1 protein might be in parallel with canonical UBA-UBL domain containing ubiquitin shuttle factor proteins (Rad23, Dsk2, Ddi1) [46], Caf1 contributes in shuttling poly-ubiquitylated proteins to 26S proteasome for degradation.

Similarly, we found that Caf1 interacts with cellular ubiquitin conjugates under native conditions and most importantly we found that Caf1 binds to recombinant linear tetra ubiquitin chains *in vitro* (Figure 6A and B). Furthermore, we discover that Caf1 associates with 19S regulatory particle complex (Figure 5B), where the docking of canonical ubiquitin shuttle factor proteins occurs at the 26S proteasome. These discoveries confirms that Caf1 is a novel ubiquitin binding protein, it shuttles ubiquitylated proteins to 26S proteasome for proteolysis by interacting with 19S regulatory particle cap.

Collectively, taken all these findings we convinced to suggest a model (Figure 8). Caf1 protein has a novel role in the UPS pathway, particularly it functions at the post-ubiquitylation step, Caf1 binds to cellular ubiquitin conjugates and 26S proteasome, thereby functions as ubiquitin shuttle factor protein and promote the proteolysis of poly-ubiquitylated proteins. The here-identified role of Caf1 as a novel ubiquitin binding protein delineate the existence of additional unidentified ubiquitin receptors in the cells. Similarly, cells lacking all the identified ubiquitin shuttle factors was shown to be viable [47]. Our attempt to identify the ubiquitin binding and proteasome interaction surface on Caf1 protein revealed that the Caf1 does not show any similarities to the reported UBA-UBL domain containing proteins, indicating that Caf1 has novel structural features that contributes to its interaction with ubiquitin and proteasome. This concept was supported by the identification of an entirely different ubiquitin interaction motif in the proteasomal subunit Dss1 protein in *S. Pombe* [48]. Hence, the discovery of such interaction surface in Caf1 protein would conceptually advance our knowledge in identifying other cellular proteins with similar binding sites. Based on our current findings and earlier reports of Caf1 in suppressing Htt103Q toxicity [23,24], we delineate that Caf1 might have widespread role in protein degradation pathway, perhaps, not restricted to UFD and N-end rule pathway substrates. Interestingly, the mammalian Caf1 was reported to complement yeast *caf1* deletion phenotype [26], demonstrating the functional conservation of Caf1 protein among eukaryotes. Therefore, it is essential to confirm if mammalian Caf1 also has role in the protein degradation pathway, such research work would identify potential Caf1 targets in the ubiquitin proteasome system (UPS) pathway.

**Figure 8:**
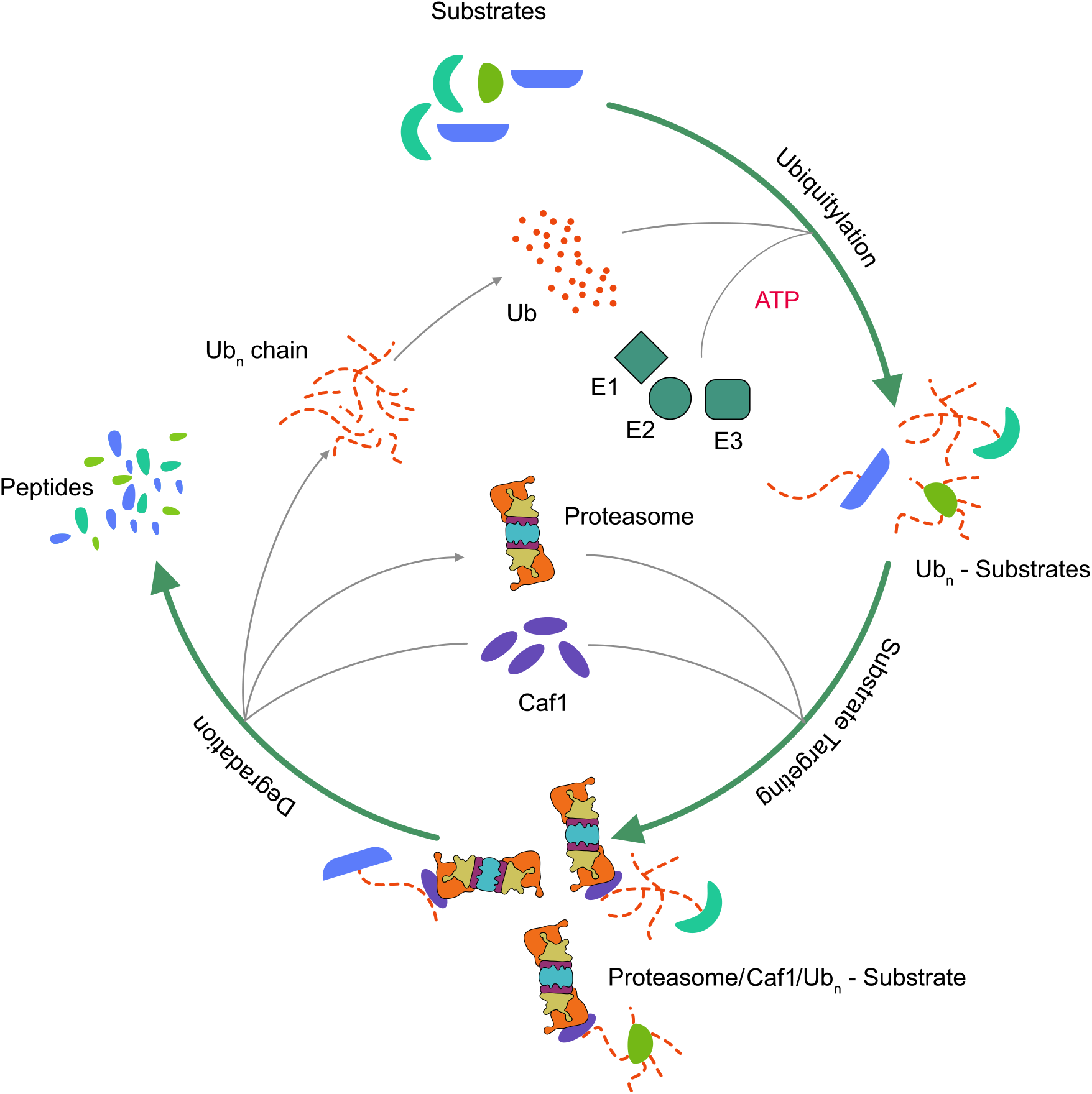
Post-ubiquitylation role of Caf1 in the UPS pathway. Model illustrates Caf1 protein in the UPS pathway. Caf1 plays ubiquitin shuttle factor role at the post-ubiquitylation step, Caf1 binds both ubiquitin-modified proteins and 19S regulatory particle subunit complex of proteasome, thereby Caf1 targets poly-ubiquitylated substrates for proteolysis by 26S proteasome.

## Acknowledgments

The presented research work was supported by funding from DBT-Ramalingaswami fellowship contingency grant as well as extra mural grant from SERB-DST to PMR. We thank Jürgen Dohmen, Marcel Fröhlich and Kerstin Nürrenberg (Institute for Genetics, University of Cologne, Germany), Ana-Mafalda Escobar-Henriques Dias and Ramona Schuster (Institute for Genetics, University of Cologne, Germany), Claes Andréasson (The Wenner-Gren Institute, Stockholm University, Sweden), Krishnaveni Mishra (University of Hyderabad, India) for their generous gifts in providing yeast strains as well as plasmids. We also extend our thanks to Sanjay Suman (CSIR-Center for Cellular and Molecular Biology, Hyderabad, India) for his help during this work. We thank Revathi Muthuvel for producing the polyclonal anti-Tpi1 antibody and Aswani Kumar for his help in preparing the graphical model used in this study.

## Supplemental information

**Supplementary figure S1:**
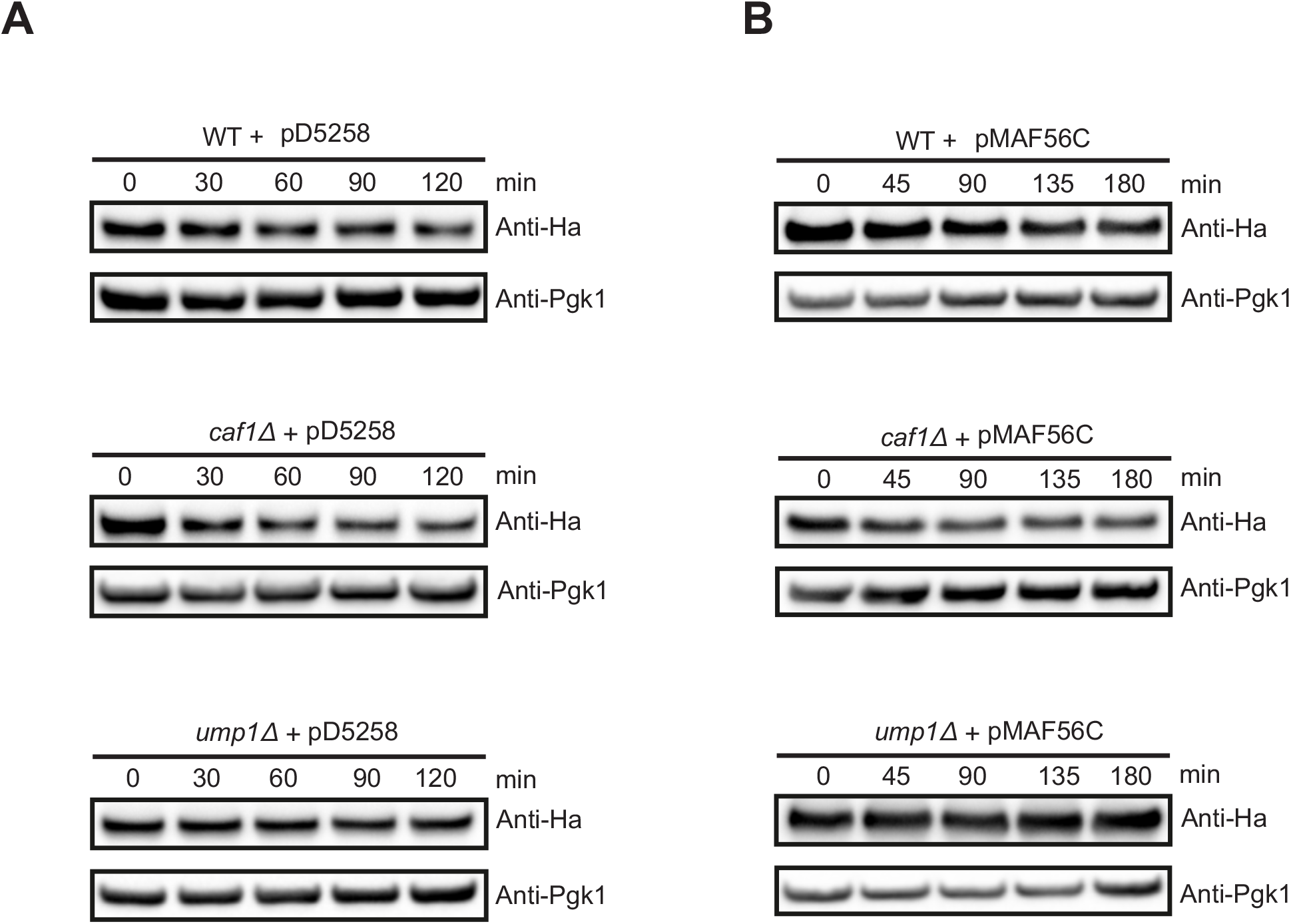
Effects of CAF1 deletion on the turn-over rates of misfolded Dhfr^mutc^ protein and ubiquitin-independent ODS-Ura3 substrate. (A) Western blot showing degradation rates of ODS-Ura3 substrate in WT, *caf1Δ* and *ump1Δ* mutant strains at the indicated time points. (B) The degradation of Dhfr^mutc^ misfolded proteins was analyzed as mentioned above. Anti-Ha antibodies were used to detect Dhfr^mutc^ and ODS-Ura3 substrate proteins. Pgk1 served as internal protein loading control.

**Supplementary figure S2:**
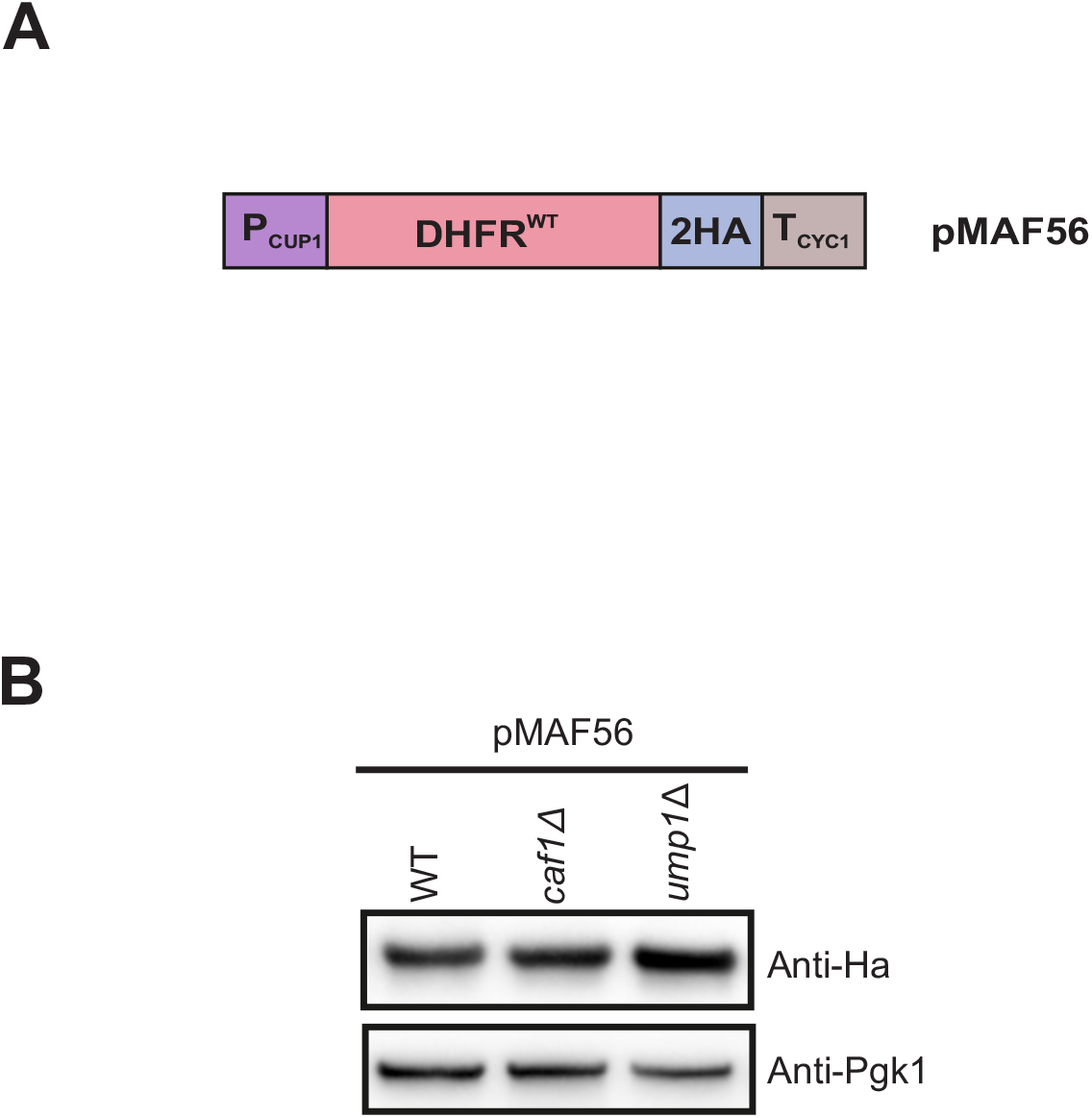
Steady state levels of stable Dhfr protein in *caf1Δ*. Western blot showing Dhfr protein levels in WT, *caf1Δ* and *ump1Δ* mutant strains at 25°C. Anti-Pgk1 and anti-Ha antibodies were used for western blotting analysis. Data represents sample volume of three independent biological samples.

